# Myristoylation and its effects on the human Golgi Reassembly and Stacking Protein 55

**DOI:** 10.1101/2021.06.22.449421

**Authors:** Emanuel Kava, Luis F. S. Mendes, Mariana R. B Batista, Antonio J. Costa-Filho

**Affiliations:** Molecular Biophysics Laboratory, Ribeirão Preto School of Philosophy, Sciences and Literature, Physics Department, University of São Paulo, Ribeirão Preto – Brazil

## Abstract

GRASP55 is a myristoylated protein localized in the medial/trans-Golgi faces and involved in the Golgi structure maintenance and the regulation of unconventional secretion pathways. It is believed that GRASP55 achieves its main functionalities in the Golgi organization by acting as a tethering factor and, when bound to the lipid bilayer, its orientation relative to the membrane surface is restricted to determine its proper *trans*-oligomerization. Despite the paramount role of myristoylation in GRASP function, the impact of such protein modification on the membrane-anchoring properties and the structural organization of GRASP remains elusive. Here, an optimized protocol for the myristoylation in *E. coli* of the membrane-anchoring domain of GRASP55 is presented. The biophysical properties of the myristoylated/non-myristoylated GRASP55 GRASP domain were characterized in a membrane-mimicking micellar environment. Although myristoylation did not cause any impact on the protein’s secondary structure, according to our circular dichroism data, it had a significant impact on the protein’s thermal stability and solubility. Electrophoresis of negatively charged liposomes incubated with the two GRASP55 constructions showed different electrophoretic mobility for the myristoylated anchored protein only, thus demonstrating that myristoylation is essential for the biological membrane anchoring. Molecular dynamics simulations were used to further explore the anchoring process in determining the restricted orientation of GRASPs in the membrane.

## 1. Introduction

The Golgi complex is a manufacturing organelle located in the heart of the exocytic pathway, responsible for transporting, modifying, and packaging proteins and lipids in eukaryotic cells (Viotti 2016). Since the 1950s, after the first observation of the Golgi apparatus under an electron microscope by Dalton and Felix (Dalton and Felix 1956), this organelle has been an extensive object of study, identified as having a central role in protein trafficking (Stalder and Gershlick 2020). The mammalian Golgi apparatus is organized as laterally linked stacks composed of flattened membranes called cisternae, ensuring a polarization relative to the endoplasmic reticulum. Such polarization is necessary for efficient protein glycosylation, maturation, and secretion(Farquhar and Palade 1998). In the 1990s, the protein GRASP65/GORASP1 (<u>G</u>olgi Re<u>A</u>ssembly and <u>S</u>tacking <u>P</u>rotein of 65 kDa) was the first member of the GRASP family to be discovered and characterized as a tethering factor involved in cisternae stacking in the *cis*-Golgi via the formation of a complex with the golgin GM130 (Barr et al. 1997; Barr et al. 1998). The second paralogue protein, GRASP55/GORASP2, was discovered two years later and was also implicated in the stacking of the Golgi cisternae(Shorter et al. 1999) through a complex with Golgin-45 in the *trans*-medial Golgi (Pfeffer 2001).

GRASPs are structurally described as formed by two main domains. The highly conserved GRASP domain (DGRASP) is myristoylated at the glycine 2 (Gly2) and anchored to the Golgi membranes (Jarvela and Linstedt 2012). DGRASP is, in turn, formed by two PDZ (<u>P</u>SD95/<u>D</u>lgA/<u>Z</u>o-1) sub-domains(Mendes et al. 2020), conventionally named PDZ 1 and 2. The interaction between GRASPs and their golgin partners involves the participation of the C-terminal region of the golgin that interacts at two distinctsites in DGRASP concurrently: the canonical peptide-binding groove of the PDZ1 and additional residues located either on the surface of the PDZ2 or in the region connecting both subdomains (Hu et al. 2015; Zhao et al. 2017). The second domain of GRASPs is the intrinsically disordered SPR (Serine and Proline-rich), which has regulatory functions and is not conserved even among evolutionary close species (Jarvela and Linstedt 2012).

The N-myristoylation at Gly2 occurs co-translationally after the cleavage of the initial methionine by methionyl aminopeptidases (Towler et al., 1987). It is catalyzed by N-terminal myristoyltransferases (NMTs), which are members of the GCN5-related N-acetyltransferases (GNAT) family (Dyda et al. 2000). In humans, the transfer of the myristate group to the N-terminal glycine is done co- or post-translationally by NMT1 and NMT2, being NMTs responsible for the myristoylation of about 2% of any proteome (Meinnel et al. 2020). Recently, mapping of myristoylation in several proteins resulted in the first description of what has been called the myristoylome (Boisson et al. 2003). The emergence of NMTs is linked to eukaryogenesis, and therefore myristoylation does not happen in prokaryotes(Meinnel et al. 2020). This lipidation contributes to regulating signaling and trafficking processes via the modulationof protein-membrane and protein-protein associations(Farazi et al. 2001). In several myristoylated proteins, anchoring is also driven by charge (McLaughlin and Aderem 1995; Sigal et al. 1994), hydrophobicity (Liu et al. 2009; Geyer et al. 1999), or triggered by ligand interaction (Borsatto et al. 2019; Olsen and Kaarsholm 2000).

The GRASP-membrane association was initially believed to be achieved through a dual mechanism involving the binding with a golgin partner and, in the great majority of cases, the N-myristoylation (Barr et al. 1997; Shorter et al. 1999). However, recent findings showed that the depletion of both mammalian GRASPs led to the loss of the golgins GM130, p115, and Golgin-45 from the Golgi, suggesting that GRASPs are responsible for the anchoring of golgins and not the opposite (Grond et al. 2020). Therefore, GRASP N-myristoylation might play an even more central role in the Golgi organization than previously believed.

The membrane anchoring has given GRASPs a multitask tethering function observed *in vitro* and in cell-based assays (Grond et al. 2020). The data reported so far indicate that tethering occurs via the *trans*-oligomerization of the PDZ domains in juxtaposed vesicles (Grond et al. 2020; Rabouille and Linstedt 2016; Feng et al. 2013). According to Truschel et al. (2011), the GRASP55 GRASP domain (DGRASP55) forms a homodimer through binding the PDZ2 of one GRASP molecule PDZ1 of the opposed protein.In a second proposed model, the homodimer is formed by a PDZ2 binding pocket-mediated interaction only (Feng et al., 2013). One major limitation in all those models of GRASP oligomerization is that they were based on the DGRASP crystal structures obtained without the N-myristoylation. Moreover, the non-myristoylated version of DGRASP55 has already been shown to behave predominantly as monomers in solution (Mendes et al. 2020; Li et al. 2013; Reddy et al. 2019; Truschel et al. 2012). Therefore, we have only a partial description of the docking and *trans*-oligomerization hitherto, and the effects of N-myristoylation have been underappreciated.

The lack of the N-myristoylation at Gly2 in previous reports was likely due to the use of recombinant DGRASPs expressed in *E. coli* and to the challenge of working with membrane-bound proteins. These points explain why, despite the known importance of myristoylation for GRASPs, the impact of this lipidation on the GRASP structural behavior has still not been properly addressed. Here, we describe an optimized protocol for the expression and purification of the myristoylated GRASP55 GRASP domain (myr-DGRASP55) in *E. coli* together with its biophysical characterization and comparison with its soluble version (DGRASP55). Our data show how myristoylation affects GRASP55 membrane anchoring tendency and structural stability with a clear impact on this protein propensity for oligomerization.

## 2. Materials and methods

### 2.1 Cloning of the HsDGRASP55 in an appropriated expression vector

The human DGRASP55 gene flanked by NcoI/XhoI restriction enzymes sites was amplified from an N-terminal 6xHis-tagged pET-28a(+)-DGRASP55 expression vector (Mendes et al. 2019) using appropriate oligonucleotides. The amplified DGRASP55 gene solution was further digested using NcoI/XhoI restriction enzymes and the product was purified using the QIAquick Gel Extraction Kit (Qiagen). The purified 621 bp fragment was ligated into the pET-28a(+) vector, previously linearized with the same restriction enzymes to form the construct. These steps were necessary to place the His-tag in the C-terminal region since the enzyme NMT from *Candida albicans* (CaNMT) requires a glycine in position 2 to perform the myristoylation. The resulting plasmids were propagated in *E. coli* DH5α, extracted with Wizard Plus SV Minipreps (Promega) and confirmed by automated DNA sequencing. The pET-28a(+)-DGRASP55 plasmids were transformed in *E. coli* Rosetta strain, grown in Luria broth (LB) medium supplemented with 50 μg/mL kanamycin and 34 μg/mL chloramphenicol. For the myristoylated protein expression, the DGRASP55 and CaNMT1-pET-22b(+) plasmids were co-transformed in *E. coli* Rosetta strain, grown in LB medium supplemented with 50 μg/mL kanamycin, 34 μg/mL chloramphenicol and 100 μg/mL ampicillin. The expression temperature was kept at 37ºC until the solution reached an optical dispersion (OD_600 nm_) of 0.8. Cells were induced by adding 0.5 mmol/L isopropyl β-D-thiogalactopyranoside (IPTG) for 18 h at 18 °C, under shaking at 220 rpm. For myr-DGRASP55 expression, myristic acid (250 μM), previously solubilized in ultrapure ethanol, was added to the cell solution together with IPTG.

### 2.2 Protein purification

Cells were harvested at 7,000xg for 10 minutes, resuspended in 20 mL of lysis buffer (20 mM Tris/HCl pH 8.0, 150 mM NaCl, 1% Triton X-100) per liter of cell culture. The cells disruption by sonication was done using a Branson 450 Digital Sonifier® (Sonitech), in ice at 48 × 5 s bursts, with an amplitude of 18% and a 15 s interval between bursts, followed by centrifugation (12,000xg, 25 min) to separate the insoluble fraction. The supernatant was loaded into a 4 mL Ni^2+^ -NTA affinity column, previously equilibrated with lysis buffer. The column was washed with 20 mL of buffer A (20 mM Tris/HCl pH 8.0, 0.03% DDM, 150 mM NaCl) and a crescent imidazole gradient (10 mM and 20 mM) used to remove weakly bound contaminants in the resin. The purified protein was eluted with 10 mL of buffer A containing 350 mM imidazole. The solution was concentrated with an Amicon Ultra-15 Centrifugal Filter (NMWL of 10 kDa, Merck Millipore, Burlington, MA, USA) and loaded into a Superdex75 10/300 GL gel filtration column (GE Health-care Life Sciences) coupled to an *Äkta purifier* system (GE Healthcare). To perform the subsequent experiments, we collected samples of DGRASP55 eluted in 16 mL to 17 mL and myr-DGRASP55 eluted from 15 mL to 17 mL. For protein concentration determination, the extinction coefficient at 280 nm was calculated with ProtParam webserver (Garg et al. 2016), resulting in ε_280_ = 26,930 M^-1^ cm^-1^ and Abs 0.1% (= 1 mg/mL) of 1.183.

### 2.3 Detergent adsorption

The removal of DDM in the protein solutions was done by fixing the protein concentration at 1 mg/ml and using 30 mg of BioBeads SM2 (Bio-Rad, CA) per mg of protein. Then, the adsorbent was incubated with purified samples for 30 minutes at 4 ºC while shaking, followed by centrifugation (13,600xg per 1 minute).

### 2.4 Circular Dichroism (CD)

CD experiments were performed in a Jasco J-815 CD Spectrometer (JASCO Corporation, Japan) equipped with a Peltier temperature control, using a quartz cell with 1 mm and 1 cm path length for far- and near-UV, respectively. The scanning speed was 50 nm·min^-1^, a spectral bandwidth of 1 nm, a response time of 0.5 s, and averaging the final spectra using 9 different accumulations. The buffer solution was 20 mM of Sodium Phosphate buffer, pH 8.0, 0.03% DDM for far-UV measurements, and a 20 mM Tris/HCl, 0.03% DDM, 150 mM NaCl, pH 8.0 for the near-UV. Protein concentration was 0.15 mg/ml for far-UV and 1.5 mg/ml for near-UV.

### 2.5 Fluorescence spectroscopy

Steady-state fluorescence was monitored using a Hitachi F-7000 spectrofluorometer equipped with a 150 W xenon arc lamp. The excitation and emission monochromators were set at a slit width of 5 nm in all experiments. The tryptophan excitation wavelength was set at 295 nm, and the emission spectra were measured from 310 up to 450 nm. Fluorescence quenching using the water-soluble acrylamide as a quencher was performed in a serial dilution from a 4 M acrylamide stock solution. The relation between the quencher concentration [Q] and the fluorescence intensity (I) was calculated through the Stern-Volmer relationship I/I_0_ = 1 + K_SV_ [Q], where I_0_ is the fluorescence intensity in the absence of the quencher and K_SV_ the Stern-Volmer constant (Gehlen 2020). The protein concentration was fixed at 15 μM, and all fluorescence assays were performed at 25°C, after 5 minutes of thermal equilibration.

### 2.6 Liposome preparation

Lipids 1,2-dimyristoyl-sn-glycero-3-phosphocholine (DMPC); 1-palmitoyl-2-oleoyl-sn-glycero-3-[phospho-rac-(1-glycerol)] (POPG); L-α-Phosphatidylethanolamine-N-(lissamine rhodamine B sulfonyl) (Ammonium Salt) (Egg-Transphosphatidylated, Chicken) (Egg LissRhodPE) were all purchased from Avanti Polar Lipids, Inc. (Alabama, U.S.A). The necessary quantity of phospholipids solubilized in chloroform was poured in glass tubes and slowly dried with nitrogen gas for liposome preparation. An additional drying step was performed using a SpeedVac™ concentrator (SAVANT™) for 2 hours. The lipid film was resuspended in buffer A without DDM, and three cycles of freeze-thaw procedure were done to disrupt multilamellar vesicles. Large unilamellar vesicles (LUVs) were prepared by submitting the freeze-thawed vesicles to an extruder with a 100 nm pore size polycarbonate membrane from Whatman (Schleicher &Schuel). The liposomes were prepared in a 4 mM stock solution supplemented with 1% Egg LissRhod PE, and the membrane composition utilized was DMPC:POPG (4:1).

### 2.7 Liposome electrophoretic mobility shift assay (LEMSA)

Liposome electrophoresis was carried in a horizontal setup, in a Tris-acetic acid solution, and the agarose gel was made in a concentration of 0.35%. The lipid samples (200 μM) were pre - incubated with protein (20 μM) and supplemented with 5% glycerol. The voltage utilized was constant (75 V), at room temperature (20 ºC – 25 ºC). The relative electrophoretic mobility of myristoylated and non-myristoylated proteins was estimated utilizing the expression: μ = d(t)/Et, where d(t) represents the distance measured from the start running point at time t, and E is the electric field used in the experiment. The image data was edited in the Adobe Fireworks CS6 software to enhance the contrast between the fluorescent bands and the background.

### 2.8 Molecular Dynamics Simulations

All-atom Molecular Dynamics simulations were performed using the crystallographic structure (3RLE) of the GRASP55 GRASP domain from *Homo sapiens* (residues 7-208). The model of the missing N-terminal was generated by homology using the Swiss model (Schwede et al. 2003) and the PDB 5GML as a template. The CHARM-GUI (Jo et al. 2008) web interface was used to generate an initial setup of the protein close to the membrane of hydrated palmitoyl-oleoyl-phosphatidylcholine (POPC) bilayer with 382 lipids units and 34,950 water molecules. To neutralize the system, Na^+^ Cl^-^ ions were explicitly added. Simulations were performed with the NAMD (Humphrey et al. 1996) package, CHARMM36 force-field, and TIP3P model for water. The PME method was used for long-range electrostatic interactions, and a cutoff of 12 Å for van der Waals forces. The isothermal-isobaric ensemble at 303.15 K, 1 atm, and a time-step of 2 fs were temperature-controlled by Langevin dynamics with 10 ps^-1^ for damping coefficient. For pressure control, the Nosé-Hoover algorithm with 200 fs of oscillation period and 100 fs for decay rate. Trajectories were visualized using the VMD package. Residue contacts (cutoff of 3 Å) and minimum distances, we utilized the software CPPTRAJ (Roe and Cheatham 2013).

## 3. Results

### 3.1 Myristoylation of GRASP55

We started by optimizing a previously described protocol used for the myristoylation of GRASP65 (Sengupta et al. 2009). We decided to change the yeast NMT1 gene used in that previous report for the well-characterized NMT cloned from *Candida albicans* (CaNMT), a stable protein with high activity and solubility (Wiegand 1992). CaNMT was cloned in a pET22b vector using the Nde1/Xho1 restriction sites, yielding a protein lacking any affinity tag. The CaNMT-pET22b (ampicillin-resistant) and the His-tagged DGRASP55-pET28a (kanamycin-resistant) were co-transformed in Rosetta (DE3). Cells were IPTG induced in the presence of 250 μM of either myristic acid or non-purified chemically synthesized azido-tagged analogues of myristic acid (Heal et al. 2002). We were not able to detect differences in the myristoylation efficiency between those reagents (data not shown). The overexpression performed for 18 h at 18 ºC also helped to increase the total amount of well-folded protein.

The success of the myristoylation protocol was indirectly checked by qualitatively evaluating the protein solubility after removing the detergent from the purified myr-DGRASP55 solution using the adsorbent Bio-Beads SM2. To do so, we expressed a version of DGRASP55 with an eGFP-tag in its C-terminus (DGRASP55/pWALDO-d). Figure 1A shows that the myristoylated protein instantly precipitated after removal of the detergent as highlighted by the green color of eGFP in the insoluble fraction after centrifugation. The control solution using the non-myristoylated DGRASP55 was shown to have its solubility utterly independent of the detergent’s presence or absence (Figure 1A). The non-polar polystyrene beads have a high surface area that adsorbs organic compounds from aqueous solutions (Rigaud et al. 1998). When caught by the Bio-Beads, the detergent micelles expose the myristoyl chain that was previously surrounded by detergent, thus leading to protein aggregation. This result suggests that the myristoyl chain in DGRASP55 was already exposed to the aqueous environment and did not seem to need a switch mechanism to bring it outwards, as previously observed for other proteins (McLaughlin and Aderem 1995; Zha et al. 2000).

**Figure 1.**
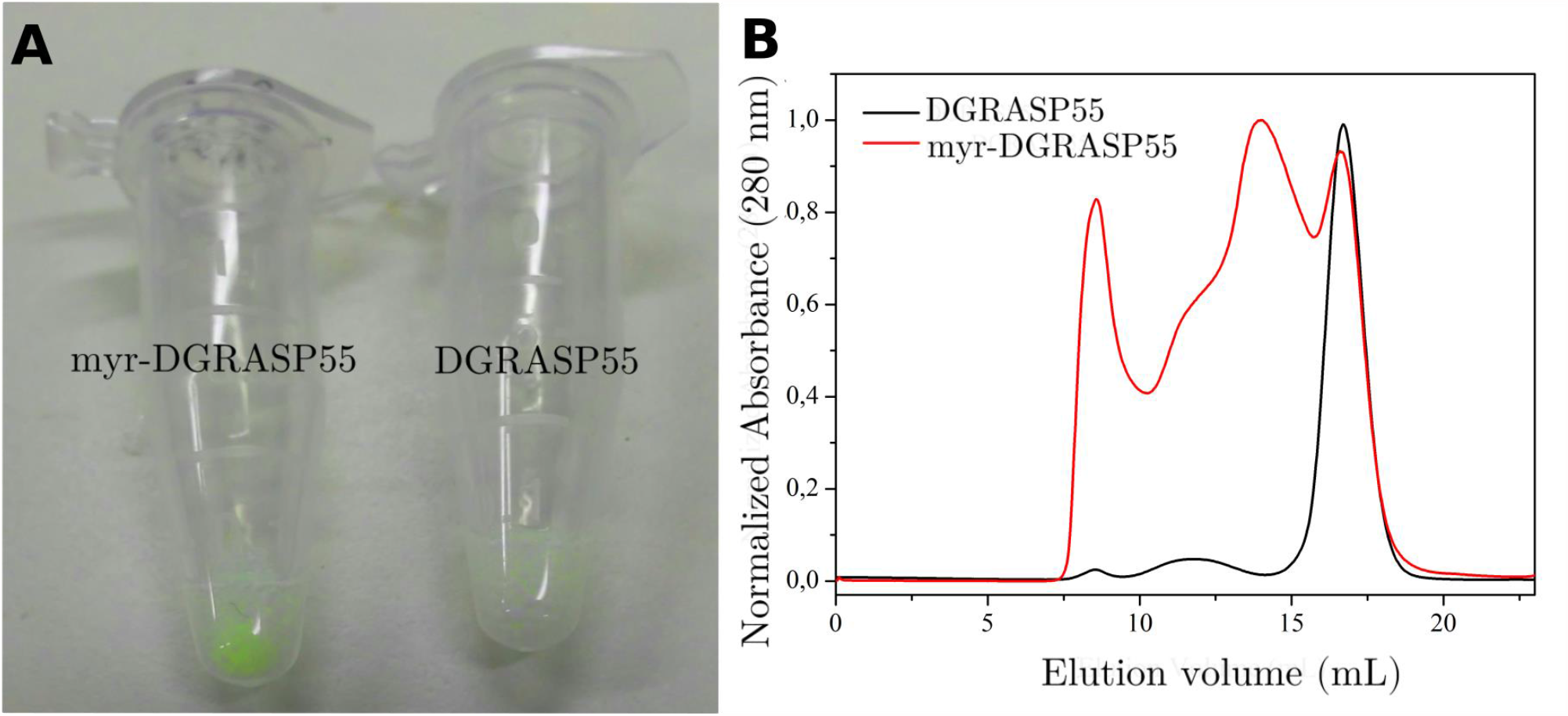
(A) Images showing the results of removing the detergent on the solubility of purified eGFP-tagged myr-DGRASP55 and DGRASP55. (B) Size exclusion chromatography data showing the myr-DGRASP55 and DGRASP55 elution profiles in detergent-containing buffer, resulting in different peaks in the elution profile of myr-DGRASP55 only.

We next monitored the protein self-association using size-exclusion chromatography (SEC) performed in the presence of the detergent. SEC data suggested that purified myr-DGRASP55 formed different oligomers with a profile that is not followed by its non-myristoylated version (Figure 1B). The DGRASP55 chromatogram was similar to those already presented in previous reports of DGRASPs (Fontana et al. 2018). The purity of the sample injected into the SEC column (See the Supporting Information, Figure S1– lane 6) was analyzed with SDS-PAGE.Since the samples were fairly pure,our SEC data indicate that the myristoyl chain is essential for GRASP oligomerization.

The success of the myristoylation protocol was also assessed by monitoring the effective anchoring of myr-DGRASP55 in lipid membranes. This was experimentally evaluated through aliposome electrophoretic mobility assay (LEMSA) after incubation of liposomes with the myristoylated and the non-myristoylated protein. A clear reduction in the electrophoretic mobility of the liposomes containing myr-DGRASP55 was observed (Figure 2). One interesting feature is the presence of a band at the top of the lane (indicated by a white arrow inFigure 2) corresponding to myr-DGRASP55-loaded liposomes that did not enter the gel. Previous data of GRASP55 tagged with a mitochondrial targeting sequence derived from the bacterial actin nucleator protein ActA of *Listeria monocytogenes* showed clustering of the mitochondria (Truschel et al. 2011). This potentially suggests that myr-DGRASP55 could induce LUV clustering by trans-oligomerization, which in principle increases the apparent size of the membrane structures, thus preventing them from permeating the agarose gel.

**Figure 2.**
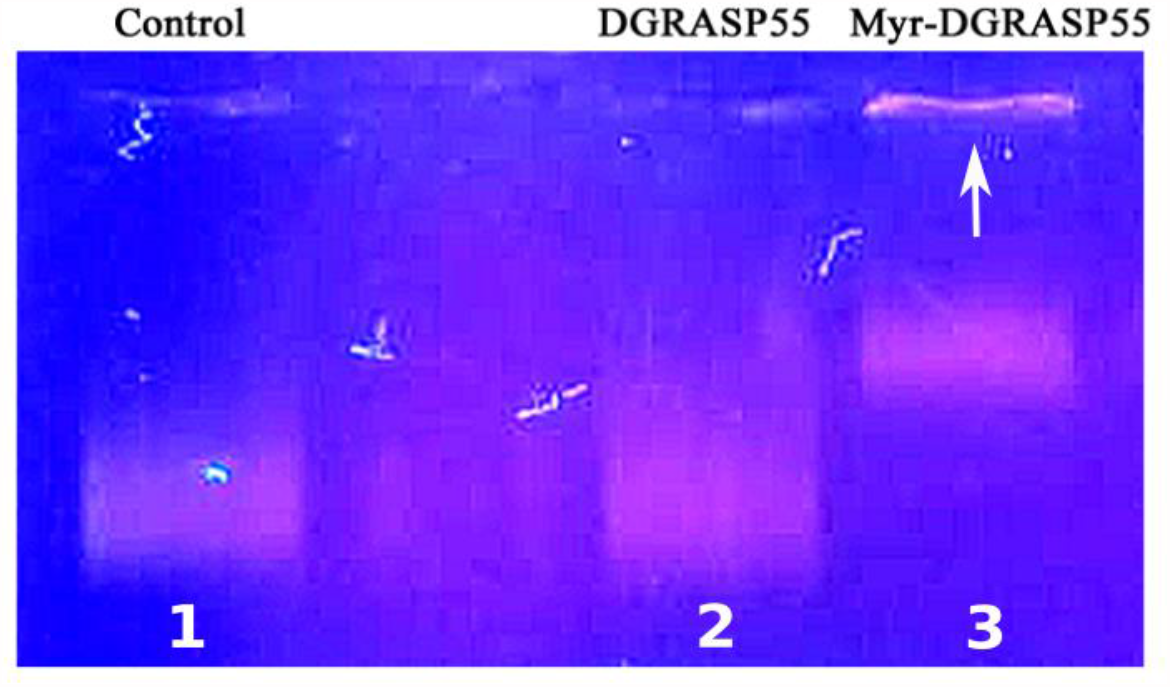
Results of the liposome electrophoretic mobility assay (LEMSA) of pre-incubated vesicles of DMPC:POPG (4:1) in agarose gel.The corresponding lanes are (1) control sample (liposomes + buffer) and samples incubated with (2) DGRASP55 and (3) myr-DGRASP55. Source: prepared by the author.

The liposomes incubated with DGRASP55 and the control sample (only liposomes in buffer solution) presented the same electrophoretic mobility of (2.59 ± 0.05) (10^−4^ cm^2^ V^-1^ s^-1^). This was significantly different from the situation observed after the incubation with myr-DGRASP55. In this case, two populations were obtained: one presenting much slower electrophoretic mobility (1.66 ± 0.05)(10^−4^ cm^2^ V^-1^ s^-1^) and a second population that appeared at the top of the respective lane in the gel (Figure 2) and could not permeate the agarose gel in the conditions tested. This indicates that myristoylation of DGRASP55 was successfully achieved, and it is essential for anchoring this protein to the membrane surface.

### 3.2 Myristoylation causes changes in the aromatic residues’ microenvironment

Once the myristoylated construction was successfully expressed, purified and the presence of the myristoyl modification demonstrated, we then moved our attention to performing a more detailed biophysical characterization of the protein under investigation. To do so, we firstly performed circular dichroism experiments to assess potential changes in the protein’s secondary structures and their spatial arrangements. The far-UV CD spectra of both proteins (Figure3) were identical, evidencing that the myristoylation did not impact the secondary structures as monitored by the ellipticity in the far-UV wavelength region (198 nm – 260 nm). The spectra followed a similar profile observed in previous reports of DGRASP55 from our group (Mendes et al. 2019).

**Figure 3.**
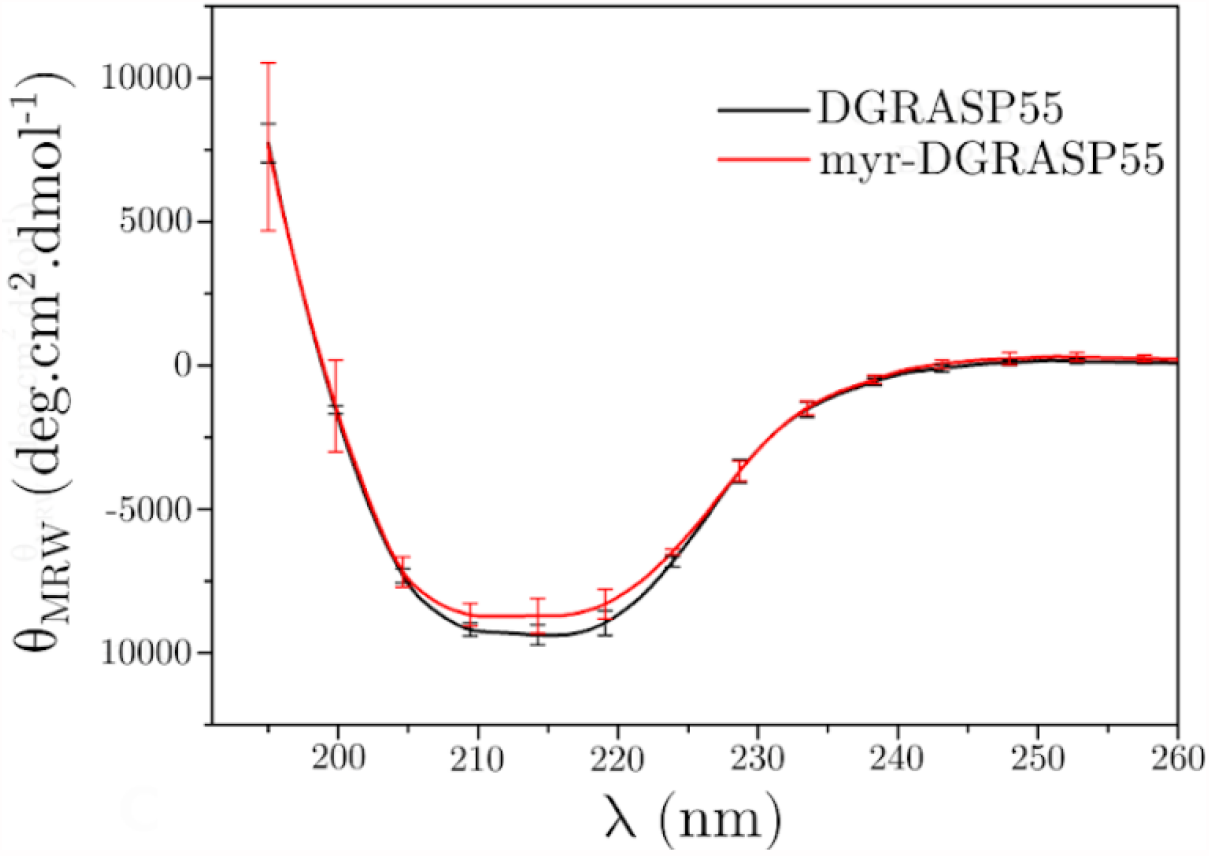
Far-UV CD spectra of DGRASP55 and myr-DGRASP55.

Although the far-UV CD spectra of the myr-DGRASP55 and DGRASP55 were indistinguishable, their thermal unfolding followed different pathways (Figure 4). While the DGRASP55 spectra transitioned from the regular far-UV CD spectrum of ordered proteins to a spectrum typical of disordered structures, the myr-DGRASP55 spectra did not present significant changes, suggesting only a slight impact on the protein secondary structure. The structural organization induced by the myristoylation of the DGRASP55 seems to give rise to a more thermal stable protein, at least in terms of maintaining its secondary structure arrangement.

**Figure 4.**
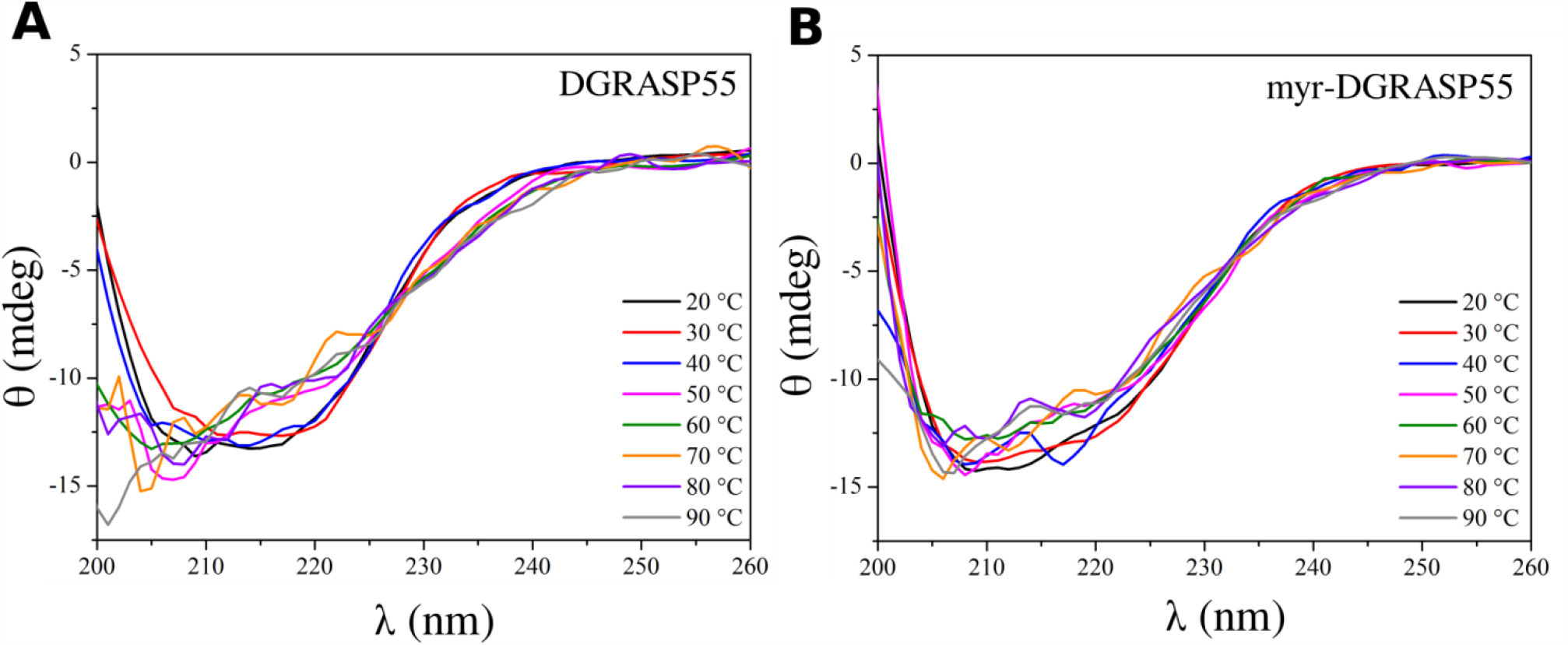
Thermal unfolding of (A) DGRASP55 and (B) myr-DGRASP55 seen by far-UV CD spectroscopy.

We again used circular dichroism experiments to explore further changes caused by the myristoylation, but now in the near-UV range. Unlike the far-UV data, the near-UV CD data showed significant differences between both versions of DGRASP55 (Figure 5A). The near-UV CD detects the optical activity of aromatic residues (Trp, Phe, and Tyr), and the intensities observed are dependent on the environment around those aromatic side chains (Li and Hirst 2017). Since there were no significant changes in the secondary structure organization monitored by the far-UV CD, the differences in the near-UV region were likely due to the alterations in the local environment of the aromatic residues in a region close to the myristoylation site.

**Figure 5.**
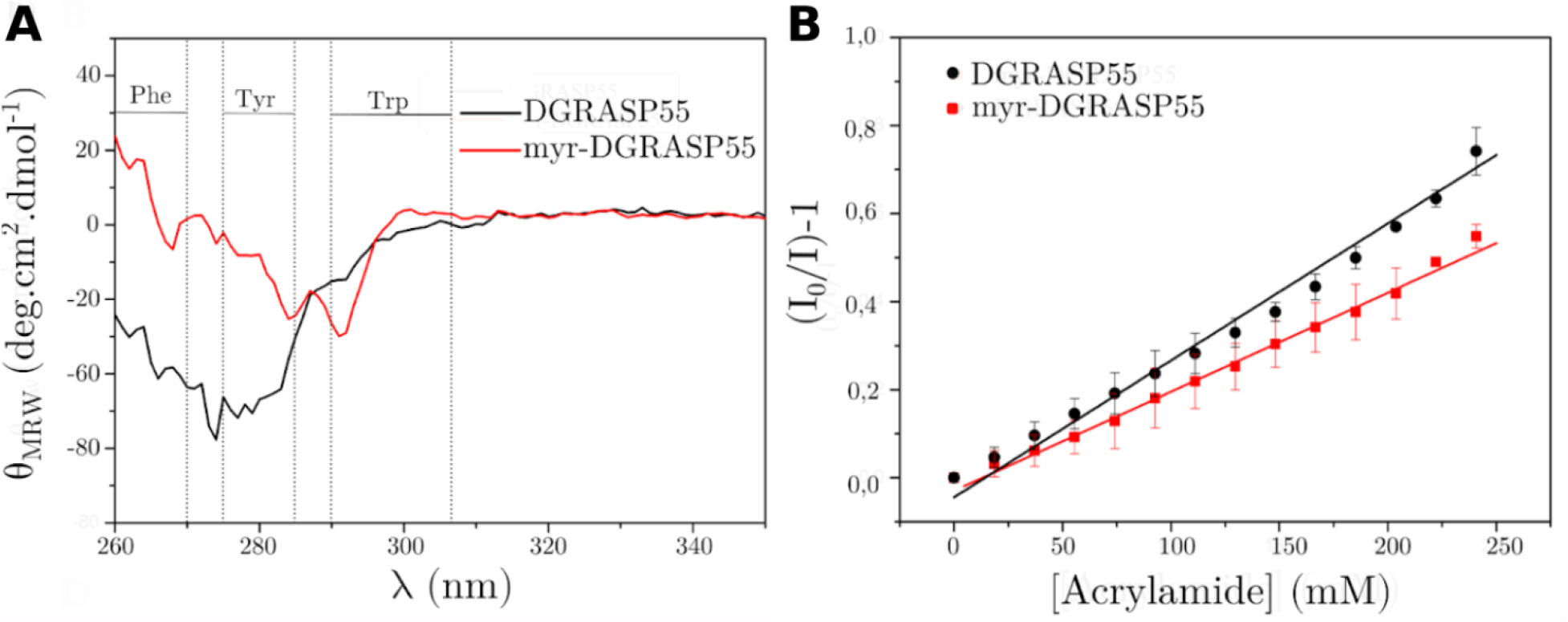
(A) Near-UV spectra of DGRASP55 and myr-DGRASP55. The dashed vertical lines delimit the regions where contributions from phenylalanine, tyrosine and tryptophan residues are expected. (C) Stern-Volmer plot showing the quenching of the tryptophan fluorescence in the presence of acrylamide. Solid lines are fits of the experimental data using the Stern-Volmer equation.

To have more information on the changes around the aromatic residues seen in the near UV-CD spectra of myr-DGRASP55, we also used steady-state fluorescence to look for specific local changes around the Trp residues. Steady-state fluorescence in the presence of the acrylamide quencher revealed that the myristoylation interfered in the accessibility of some tryptophan residues (Figure 5B). The Stern-Volmer constant (K_sv_) of 2.25 M^-1^ obtained for the myristoylated protein compared with K_sv_ = 3.11 M^-1^ obtained for DGRASP55 indicates that the tryptophan residues are less quenched (more protected from the acrylamide) in the myristoylated protein. This result is supported by the differences in aromatic residues environment in the near-UV CD data. It suggests changes in the microenvironment of at least one tryptophan residue of DGRASP55 after myristoylation and binding to the detergent molecules. These observations are discussed in detail in the MD simulations section below, where the surroundings of the aromatic residues in DGRASP55 and membrane-anchored myr-DGRASP55 were explored. Therefore, our near-UV and fluorescence quenching data suggest that some of the myr-DGRASP55 tryptophan residues are located in a different local environment upon myristoylation and solubilization with DDM micelles.

### 3.3 Exploring the DGRASP-membrane interaction through MD simulations

Previously, Heinrich *et al*. (2014) reported neutron scattering data on the orientation of DGRASP55 in a membrane model system. In that case, the anchoring to the membrane was achieved using both the Gly2 myristoylation and the interaction of a 6 -His-tag located in the C-terminus of DGRASP55 with a 1,2-dioleoyl-sn-glycero-3-[(N-(5-amino-1-carboxypentyl)iminodiacetic acid)succinyl] phospholipid inserted in the model membrane. Therefore, our simulations of the DGRASP55 started with the protein in an orientation relative to the membrane as described by Heinrich *et al*. (2014) (Figure 6A). Moreover, the simulations of myr-DGRASP55 started in a configuration where the myristoyl chain was partially inserted into the bilayer (Figure 6B), resulting in the restriction of the protein to a position close to the membrane, which was maintained along all calculated trajectories (Figure 6C).

**Figure 6.**
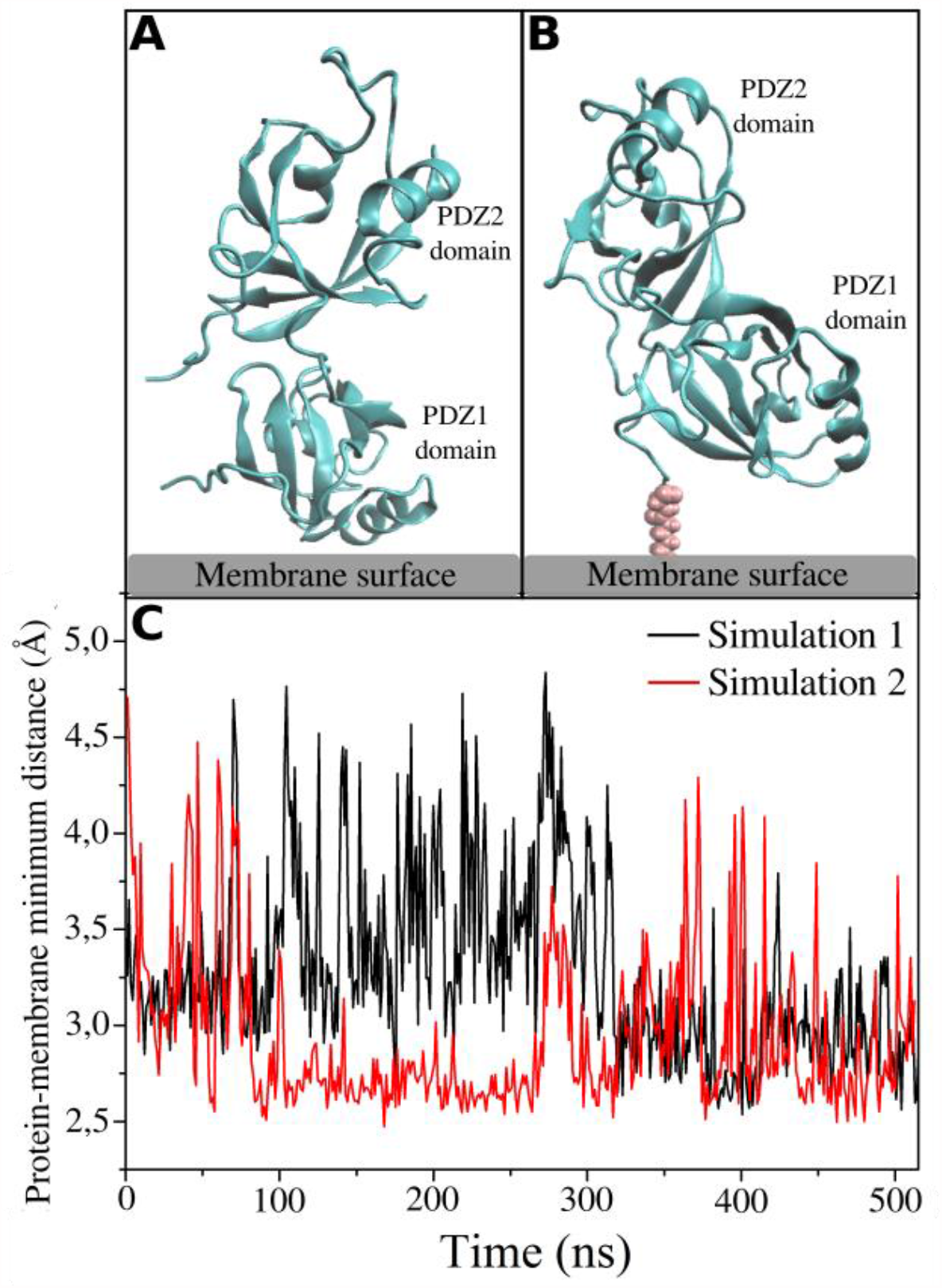
(A) DGRASP55 initial orientation relative to the membrane surface described by Heinrich et al. (2014). (B) myr-DGRASP55 initial orientation used in our simulations. (C) Minimum distance between the atoms of anchored myr-DGRASP55 and the lipid residues in two different trajectories. After around 300 ns (Simulation 1) and 100 ns (Simulation 2), the myristoyl chain penetrated slightly more into the bilayer, and the minimum distance was stabilized.

In our simulations of DGRASP55, we observed the protein detaching from the membrane (Figure 7A-B) in a short timescale of approximately 800 ns, thus highlighting the instability of any interaction between the non-myristoylated protein and the lipid membranes. This agrees with our experimental results in Figure 1. In the simulations of myr-DGRASP55 (Figure 7C), the membrane anchoring was achieved without the need for the artificial 6-His-tag anchoring strategy used before, which likely resulted in a tilted orientation towards the membrane surface in our model (Figure 7D), when compared to that used by those authors (Heinrich et al. 2014). As a result, some of the aromatic residues of myr-DGRASP55 (highlighted in colors in Figure 7C-D) were localized close to the interface between the protein and the membrane. As said above, we did not observe the detachment of the protein due to the myristoyl-anchoring, which kept those aromatic residues (Tyr71, Trp89, and Phe206)close to the interfacial region. Furthermore, we observed a series of residues that were in constant contact (distance < 3.0 Å) with the phospholipids along the simulated trajectories(Figure 8). The residues in contact with the bilayer belong to the following regions: residues 43 to 45 (Asn43, Gly44, and Ser45), 60 to 67 (Ala60, Asn61, Val62, Glu63, Lys64, Pro65, Val66, and Lys67) and 80 to 82 (Glu80 and Ser82) (Figure 8B-C). The presence of this network of interactions is important because it was shown before that the binding energy provided by the myristoylation only is weak (K_D_ of ∼10^−4^ M), and hence insufficient to fully anchor a protein to the membrane (Resh 1999). Our MD data suggest that a coupled mechanism, which includes the Gly2 myristoylation and a series of interactions between amino acids of myr-DGRASP55 with the membrane surface, is responsible for the restricted configuration of GRASP55 in membranes, necessary to avoid the *cis*-oligomerization.

**Figure 7.**
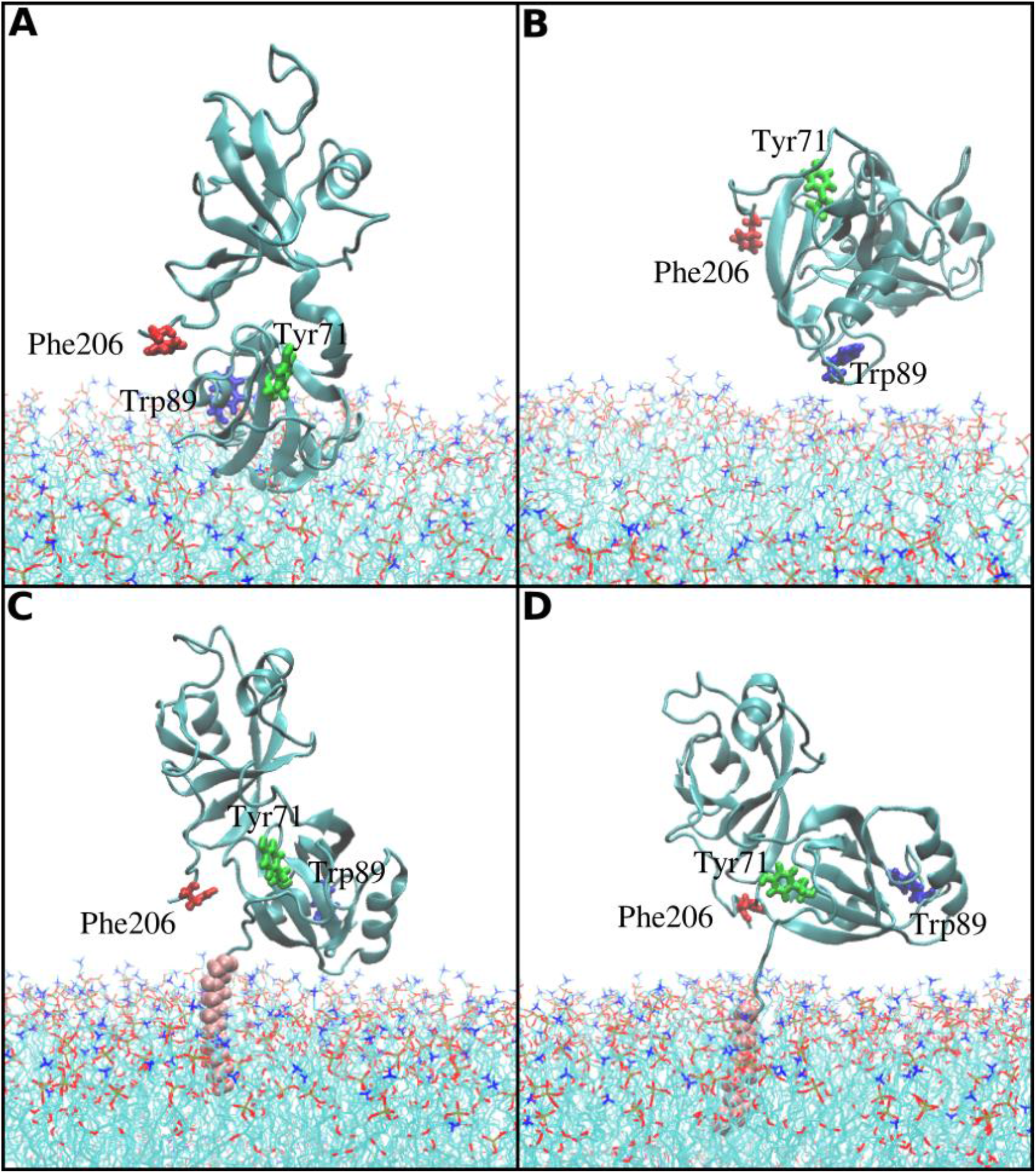
MD simulations of DGRASP55 and myr-DGRASP55 in contact with a model membrane. Mobility of DGRASP55, starting from the orientation represented in (A) and resulting in the detachment from the membrane surface observed in (B) after 1.4 μs. Simulations of the myr-anchored protein started with the orientation represented in (C) and resulting in the orientation shown in (D) after 780 ns. The aromatic residues are shown in licorice representation: Trp89 (blue), Tyr71 (green) and Phe206 (red). Pink spheres represent the myristoyl group. Lipids in the membrane are shown as lines, hydrocarbon chains are colored in cyan, and headgroups are colored in red (oxygen), ocher (phosphorus), and blue (nitrogen).

**Figure 8.**
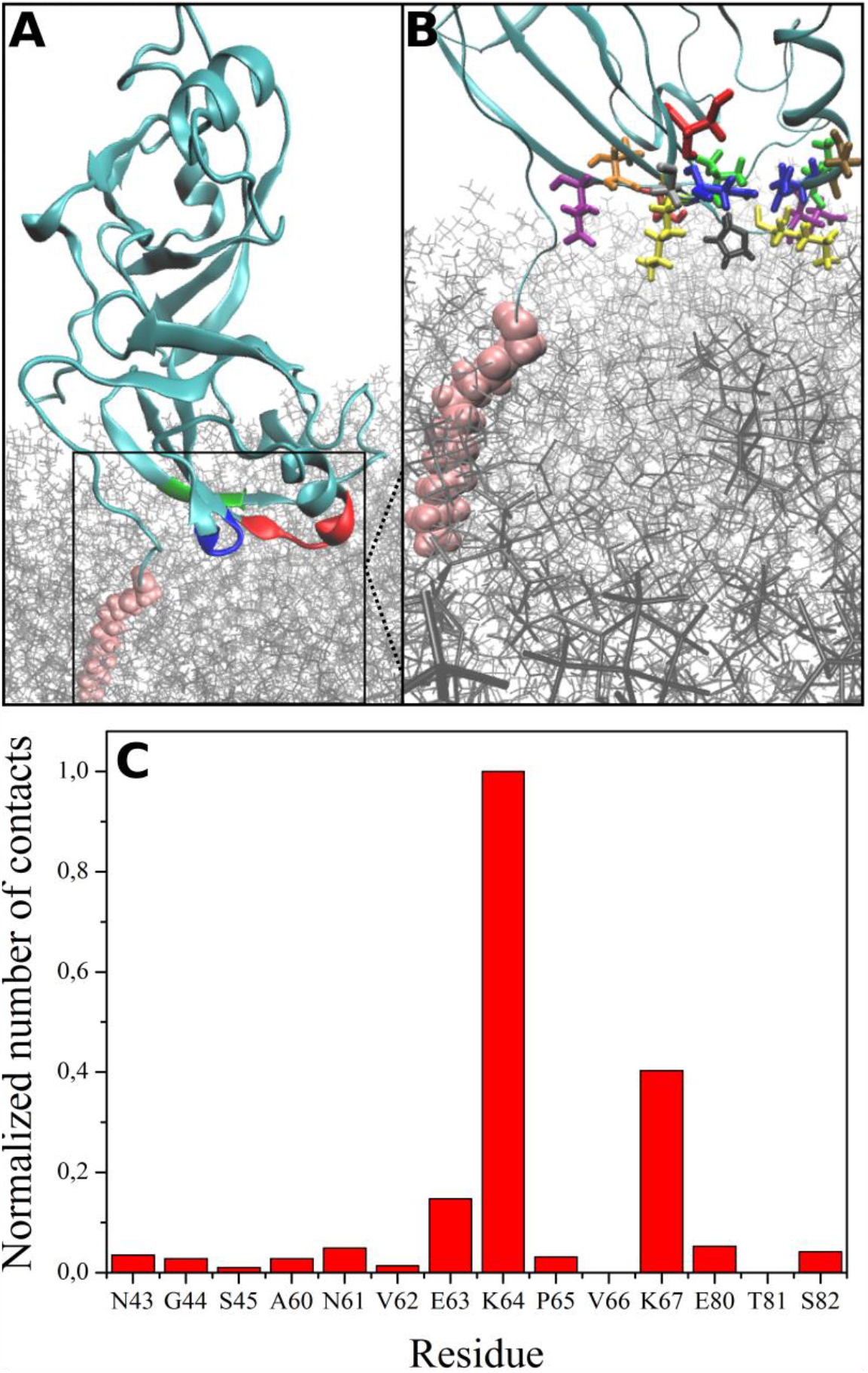
(A) Regions at the protein-membrane interface in contact with the lipid bilayer. The regions are colored in blue (residues 43 to 45), red (residues 60 to 67), and green (80 to 82). (B) Details of the interfacial region identifying specific amino acid residues, drawn as licorice representation and colored in blue (Asn43 and Asn61), red (Ser45 and Ser82), yellow (Lys64 and Lys67), gray (Gly44), green (Val62 and Val67), orange (Thr81), purple (Glu63 and Glu80), ocher (Ala60) and black (Pro65). Lipids are represented as gray sticks, and myristate is represented as pink spheres.

## 4. Discussion

GRASPs are peripheral membrane proteins, whose mechanism of anchoring, despite still lacking a detailed molecular understanding, has been proposed in general terms. Such mechanism, in most GRASPs, was suggested to involve the myristoylation of Gly2 of the GRASP domain along with the participation of a partner protein (Rabouille and Linstedt 2016). The crystal structures obtained so far for the GRASP domains (Feng et al. 2013; Truschel et al. 2011) were determined using protein constructions that lacked the myristoylation. Therefore, the description of this lipidation in the anchoring process has been somewhat limited. Due to its relevance in the functional cycle of GRASPs, it is still an issue that deserves more attention for a better understanding of the extensively discussed GRASP oligomerization (Rabouille and Linstedt 2016).

Despite the plasticity of GRASPs in terms of structural organization, which includes the formation of dimers and fibrillar structures (Fontana et al. 2018; Reddy et al. 2020), DGRASPs in solution have shown to be predominantly monomers (Li et al. 2013; Truschel et al. 2012). Furthermore, neutron reflection experiments suggested a restriction of the myr-DGRASP55 anchored in lipid membranes, affecting the potential of myr-DGRASP55 for self-interaction. Here, we explored the effects of myristoylation in one of the human GRASPs (GRASP55) and how this post-translational modification affects the protein structural behavior and its interaction with lipid membranes.

A successful myristoylation strategy that can be implemented during the heterologous expression of the protein in bacteria is a crucial step to obtain information on GRASP in scenarios that are more closely related to those found in the cell. Our myristoylation protocol was adapted from previous reports (Sengupta et al. 2009; Heinrich et al. 2014). The success in lipidating the DGRASP55 was assessed by different methods, as seen in Figure 2 and Figure 2. Several myristoylated proteins were already shown to utilize a controlled mechanism for binding to the membranes. These mechanisms can be represented by the interaction between hydrophobic residues, negatively charged residues, and co- and post-translational modifications (Whited and Johs 2015). For example, the Golgi-localized ARF1 (ADP ribosylation factor-1) requires the presence of GTP for membrane binding (Liu et al. 2009), and Recoverin relies on a calcium-mediated conformational change to expose its myristoyl moiety (Borsatto et al. 2019; Timr et al. 2017). Our results (Figure 1 and Figure 2) indicate that DGRASP55 has its myristoyl group already exposed to the solvent, therefore requiring the use of detergent to solubilize the myristoylated protein.

Once the myristoylation of DGRASP55 was successfully achieved, we further explored the effects of this post-translational modification on the protein’s biophysical properties using a combination of experimental and computational methods. Our CD data measured in the far-UV range (Figure3) showed that the presence of the myristoyl moiety and the detergent molecules resulted in no changes to the overall protein’s secondary structure, which suggests DGRASP55 would maintain its spatial organization upon myristoylation. On the other hand, alterations in the vicinity of the aromatic residues were seen in our near-UV CD data (Figure 5A), which indicates that local rearrangements would be expected upon interaction with the Golgi or other functionally relevant membranes. Another significant alteration, introduced by the myristoyl group and the detergent micelles, was seen in the thermal unfolding of DGRASP55 (Figure 4), which yielded different behavior upon temperature increasing. Myristoylation of proteins where the myristoyl is involved in protein-protein interactions showed increased stability (Geyer et al. 1999; Sowadski et al. 1996; Muralidhar et al. 2005). For example, the myristoylation of human insulin induces structural effects that result in stable hexamers (Olsen and Kaarsholm 2000). Myristoylation can also increase protein stability by inserting the lipid moiety into a hydrophobic pocket, such as observed in forming a recognition motif in the phosphorylation site of protein kinase A (Sowadski et al. 1996).

On the other hand, myristoylated proteins showed a decrease in solubility (Geyer et al. 1999; Berthiaume and Resh 1995), and in enthalpy of unfolding (Schroder et al. 2011) compared with its non-myristoylated version. In the case of myr-DGRASP55 and DGRASP55, the presence of the myristoyl chain and the detergent yielded different thermal unfolding profiles (Figure 4). The myr-DGRASP55 protein had its far-UV CD spectrum unchanged as the temperature was raised, suggesting that myristoylation increased DGRASP55 stability.

The aforementioned changes in the local environment around the aromatic residues were also assessed by fluorescence and molecular dynamics simulations. More specifically, using steady-state fluorescence, the accessibility of the Trp residues present in DGRASP55 to a water-soluble quencher was measured. The lower Stern-Volmer constant obtained for myr-DGRASP55, compared to its non-myristoylated form (Figure 5B), can be partially explained by the proximity of Trp89 to the lipid bilayer (Figure 7D). In the non-myristoylated protein, the movement of that residue was not restricted by the myristoyl anchoring to the membrane, thus keeping it readily accessible to the quenching agent. In the experimental conditions, the solubilization of the myristoyl moiety by the detergent molecules that reduced quencher accessibility likely played a similar role to the anchoring in the lipid bilayer, which would also reduce Trp89 exposure due to its proximity to the membrane surface. The differences in near-UV CD data (Figure 5A) observed in the comparison of both protein versions can be explained by the differences in the microenvironment of the aromatic residues (Tyr71, Trp89, Phe206) shown in Figure 7D. These differences arose because the residues located at the interface of the myristoyl region with the membrane-mimicking micellar surface are less water-accessible when compared with the same positions in the DGRASP55, which does not seem to interact with the lipid bilayer as inferred by our LEMSA data (Figure 2) and MD data (Figure 7). Moreover, we identified specific regions containing residues in contact (distance < 0.3 nm) with the lipid bilayer surface (Figure 8). The higher frequency of charged residues contacts was a characteristic also observed in our MD data. Based on the residues network in contact with phospholipids, we suggest that the final protein orientation in the simulations is favored by electrostatic interactions involving specific regions (Figure 8A-B). A recent study using a DNA-based voltmeter found a high resting membrane potential measured at the trans-Golgi network, with a positively charged lumen (Saminathan et al. 2021). Despite the dynamics of the Golgi membrane surface potential depends on several factors, for example, the influence of Na^+^/K^+^ ATPases, the charged membrane surface possibly impacts the interaction between phospholipids and residues mentioned above (regions 43 to 45 (Asn43, Gly44 and Ser45), 60 to 67 (Ala60, Asn61, Val62, Glu63, Lys64, Pro65, and Lys67) and 80 to 82 (Glu80 and Ser82)) (Figure 8C).

In *in vivo* studies, GRASP55 was found in the endoplasmic reticulum under cell stress (Kim et al. 2016). This relocalization of an anchored protein likely required a membrane dissociation that would involve the myristoylated N-terminal region. Although our data indicated the solvent exposure of the myristoyl moiety, the variety of specific interactions of the full-length myr-GRASP55 could still probably include the need for a myristoyl switch, a feature observed in several myristoylated proteins as a mechanism to expose the initially sequestered myristoyl chain upon the onset of specific conditions (MARCKS (McLaughlin and Aderem 1995), Recoverin (Borsatto et al. 2019; Timr et al. 2017), ARF1 (Liu et al. 2009)). Finally, GRASPs are the most phosphorylated Golgi proteins during mitosis (Wang et al. 2003), resulting in GRASP dimers disassembly and cisternae unstacking. This indicates the relevance of including phosphorylation in future studies involving myristoylated GRASPs and its role in the transition of the protein from the Golgi to other cell locations.

## 5. Conclusions

The biophysical properties of myr-DGRASP55 are significantly different compared to the non-myristoylated protein. The myristoylation rendered a protein that is less soluble and more thermally stable. On the other hand, an explanation for the differences observed between these proteins in size-exclusion chromatography might be a consequence of protein self-association. Although there were no detectable differences in the secondary structure organization observed by the far-UV CD spectra, the lipidation altered the unfolding process monitored via the temperature dependence of the far-UV CD spectra. Additionally, we searched for a more detailed description of the membrane interaction mechanisms through molecular dynamics simulations and lipid electrophoretic mobility assays, confirming that myristoylation is essential for GRASP domain anchoring into lipid membranes or detergent micelles. Furthermore, we found that a coupled mechanism of binding is responsible for the restricted configuration of GRASP55 in membranes, and this is achieved without the need of a golgin partner, or a mimetic of it. Since myristoylation is essential for GRASP’s attachment to the membrane, these results present new insights into GRASP-membrane interaction and the effects of lipidation on the biophysical properties of these proteins.

## Acknowledgements

The authorsthankthe Fundação de Amparo à Pesquisa do Estado de São Paulo (FAPESP) (Grants No. 2015/50366-7 and 2012/20367-3), Conselho Nacional de Desenvolvimento Científico e Tecnológico (CNPq) (Grant no. 306682/2018-4) and CAPES for the financial support. EK thanks CAPES for a scholarship (88882.378790/2019-01), LFSM and MRBB thanks FAPESP for the postdoctoral grants No. 2017/24669-8 and No. 2016/16328-3, respectively. We are grateful to professor Giuliano Clososki for sharing the undecanoic acid.

## Conflicts of interest

The authors declare no conflict of interest.

## Research involving Human Participants and/or Animals

Our research does not involve human or animal participants.

## Consent to publish

All authors have consented to publish the data.

## Notes

### Competing Interest Statement

The authors have declared no competing interest.

