## Supplementary figure for "Myristoylation and its effects on the human Golgi Reassembly and Stacking Protein 55"

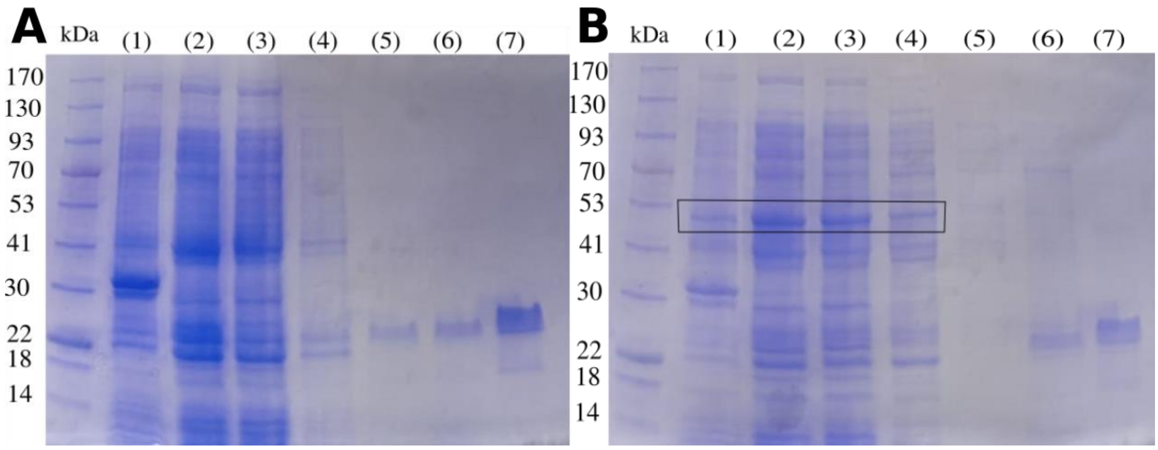


Figure S1 - (A) DGRASP55 and (B) myr-DGRASP55 purification in $Ni^{2+}$-NTA resin, visualized through SDS-PAGE; corresponding bands to CaNMT1 are highlighted by the black rectangle: (1) pellet after lysed cells centrifugation, (2) supernatant (lysate), (3) void (not bound to the column), (4) 20 mL washing buffer (Tris 20 mM, NaCl 150 mM, β-Mercaptoethanol 5 mM, DDM (0,03%) + imidazole 10 mM, (5) 20 mL washing buffer + imidazole 20 mM, (6) protein elution with 10 mL buffer + imidazole 350 mM, (7) fraction collected after gel filtration purification (Superdex200). Source: prepared by the author.
